# Detection of attomolar concentration of heart-type fatty acid binding protein using ion current rectification sensing with conical SiO_2_ nanopores

**DOI:** 10.64898/2026.04.07.717075

**Authors:** Nahid Afrin, Shankar Dutt, Maria Eugenia Toimil-Molares, Patrick Kluth

## Abstract

Rapid and highly selective sensing of ultra-low concentration protein biomarkers remains a critical challenge important for early disease diagnosis and monitoring. Here, we use conical SiO_2_ nanopore-based biosensing for the rapid detection of heart-type fatty acid binding protein (H-FABP). Antibodies were covalently immobilized on the nanopore surface through siloxane chemistry. The functionalized asymmetric nanopores generate a characteristic rectifying current–voltage response, which shows a distinct shift upon binding to the target protein due to partial neutralization of the negatively charged pore surface. The sensor exhibits excellent sensitivity in the attomolar to nanomolar concentration range with a detection limit (LOD) of ∼0.4 aM. Furthermore, the platform exhibits high selectivity, distinguishing H-FABP from non-target proteins (HSA and Hb) at concentrations six orders of magnitude higher. We also demonstrate that nanopores can be regenerated using sodium hypochloride and O_2_ plasma treatment, enabling repeated functionalization and reuse.

## Introduction

The real-time detection of disease-related biomarkers in blood or saliva at ultra-low concentrations is highly desirable for early diagnosis and point-of-care testing. Many clinically relevant biomarkers are present in biological fluids at concentrations far below the detection limits of conventional assays, particularly during the early stages of disease.^1^ While established analytical techniques such as enzyme-linked immunosorbent assays (ELISA) and electrochemiluminescence immunoassays (ECLISA) offer high specificity, they require complex instrumentation and extended analysis times, limiting their suitability for decentralized, low cost and quick diagnostics.^2^ These limitations have driven the development of new portable biosensor technologies capable of detecting biomarkers at extremely low concentrations in real time.

Solid-state nanopores provide a powerful platform for biomolecular detection by directly transducing molecular interactions into ionic current signals.^3,4^ Ion current rectification (ICR) sensing has emerged as a promising nanopore sensing technique that has enabled versatile detection of ions, small molecules, nucleic acids, and proteins.^5^ It utilizes nanopores with asymmetric geometries that exhibit ICR governed by the pore geometry and surface charge.^6^ When specific probe–analyte interactions within the nanopore alter the surface charge, measurable changes in current-voltage characteristics can be used as a measure of the presence and concentration of the analyte.^7^ However, fabrication of suitable asymmetric nanopore systems with desired geometry and surface properties remains a challenge. We have recently developed conical nanopores in SiO_2_ membranes that are particularly attractive for ICR sensing due to their controllable geometry, chemical robustness, and well-established surface chemistry. ^8,9^ The conical geometry yields strong current rectification, while SiO_2_ surface enables versatile functionalization through silane chemistry and covalent biomolecule attachment. Moreover, multipore membranes with tens to thousands of pores can be fabricated and provide ensemble-averaged electrical responses which improve stability and reproducibility compared to single-pore devices. ^10^

To demonstrate the capabilities of our nanopore membranes for biosensing, in this work we present the detection of heart-type fatty acid binding protein (H-FABP, also known as FABP3), a biomarker with clinical relevance and diagnostic versatility. H-FABP is a well-established biomarker of cardiac injury, enabling early differentiation between unstable angina and acute myocardial infarction, as it is rapidly released into the bloodstream and can be detected within approximately 1.5 hours after the onset of symptoms.^11–14^ Its diagnostic utility has led to recommendations for its use alongside cardiac troponins to improve clinical sensitivity. ^15,16^ Beyond cardiovascular applications, elevated H-FABP levels have also been reported in cerebrospinal fluid from patients with neurodegenerative disorders, including Alzheimer’s disease and synucleinopathies. ^17^ However, detection in peripheral blood remains challenging due to dilution across the blood–brain barrier requiring ultra-sensitive quantification techniques.^17^

Recent nanotechnology-enabled H-FABP biosensors, including electrochemical immunosensors, molecularly imprinted polymer sensors, capacitive immunosensors, and microcantilever-based devices, typically rely on signal amplification strategies such as redox mediators, thermal transduction, or nanomaterial-assisted electrochemical readout, achieving limits of detection in the ng/mL range (picomolar concentrations).^12,18–20^ In contrast, the sensing platform presented in this study provides direct electrical detection with attomolar sensitivity.

## Results and discussion

The nanopore membranes used in this experiment were fabricated in SiO_2_ using the track etching technique shown schematically in figure 1a.

**Figure 1:**
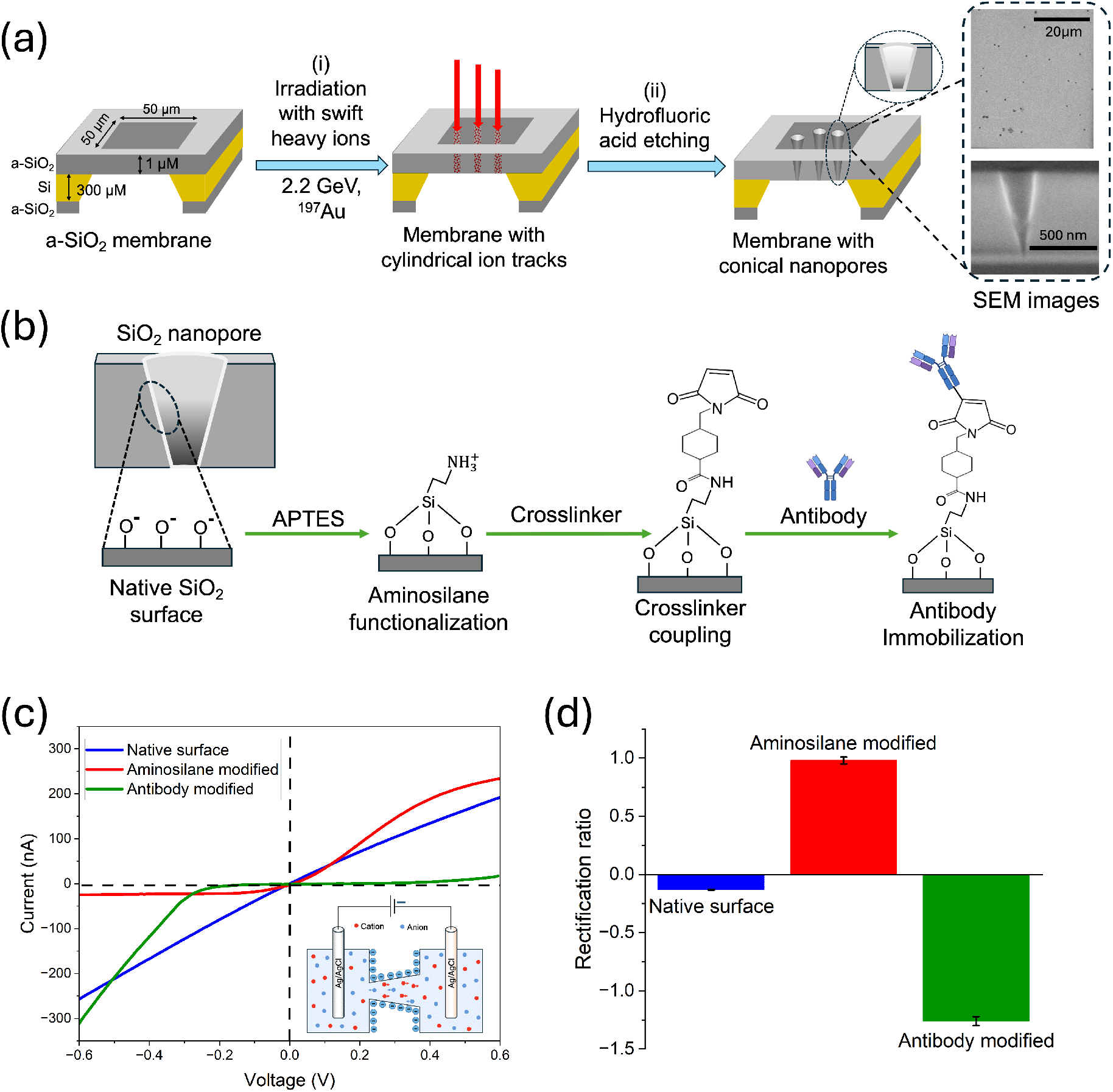
SiO_2_ nanopore fabrication and functionalization. (a) Schematic of the nanopore fabrication process, including membrane design, ion irradiation, and asymmetric track etching to form conical nanopores. (b) Schematic of the stepwise surface modification strategy: native SiO_2_ nanopore, aminosilane functionalization followed by covalent immobilization of antibodies on the pore walls. (c) Representative current-voltage (I-V) characteristics recorded after each functionalization step. (d) Corresponding rectification coefficients calculated from the current-voltage curves reflect the changes in surface charge during each stage of functionalization. The native SiO_2_ surface displays a negative rectification, whereas aminosilane modification results in a positive ratio. The rectification ratio becomes strongly negative after the attachment of highly negatively charged antibody molecules on the nanopore surface. Error bars indicate the standard error of the mean obtained from three independent measurements (N = 3) with the same nanopore membrane.

Ion-tracks were formed by irradiation of 50 × 50 *µ*m^2^ free standing SiO_2_ windows supported by Si frames with 2.2 GeV ^197^Au ions at a fluence of 1 × 10^6^ ions cm^*−*2^. Subsequent one-sided wet chemical etching in 2.5% HF leads to the formation of conical pores due to the higher susceptibility of the ion-tracks to chemical etching compared to the surrounding material. The track etching technique enables precise control over the nanopore geometry and extremely narrow size distributions, which are critical for achieving reliable and high-performance nanopore-based biosensing. A detailed description of the pore formation process is available in our previous works.^8,9^ Plan-view and cross-sectional scanning electron microscope (SEM) images were used to determine the pore density, pore geometry, and base diameter of the nanopores (see Figure 1a and Supporting Information, Figure S1a–e). From the SEM images, the pore density was estimated to be 1 × 10^6^ pores cm^*−*2^, corresponding to ∼25 nanopores within a 50 × 50 *µ*m^2^ membrane window, with an average base diameter of (380 ± 5) nm. For each membrane used in the experiments, the number of pores was individually determined from SEM image analysis. Cross-sectional SEM confirms the conical shape of the pores. The pore length was determined to be (690 ± 5) nm using ellipsometry (JA Woollam M-200D) by measuring the membrane thickness after etching. The nanopore cone angle and the pore size distribution were characterized by small-angle X-ray scattering (SAXS). The one-dimensional fitting of the SAXS data yielded a narrow size distribution of 2-4%. A detailed description of the characterization procedures is provided in our previous work.^8,21,22^ The nanopore tip diameter of the membrane containing 25 nanopores and reported in Figures 1 and 2 was estimated to be approximately 67 nm based on conductometric measurements, as described in detail in our previous work.^8,9^ Figures 3 and 4 correspond to measurements performed on independently fabricated membranes under similar conditions, containing 23 and 43 nanopores, respectively, with estimated nanopore tip diameters of approximately 69 nm and 48 nm.

**Figure 2:**
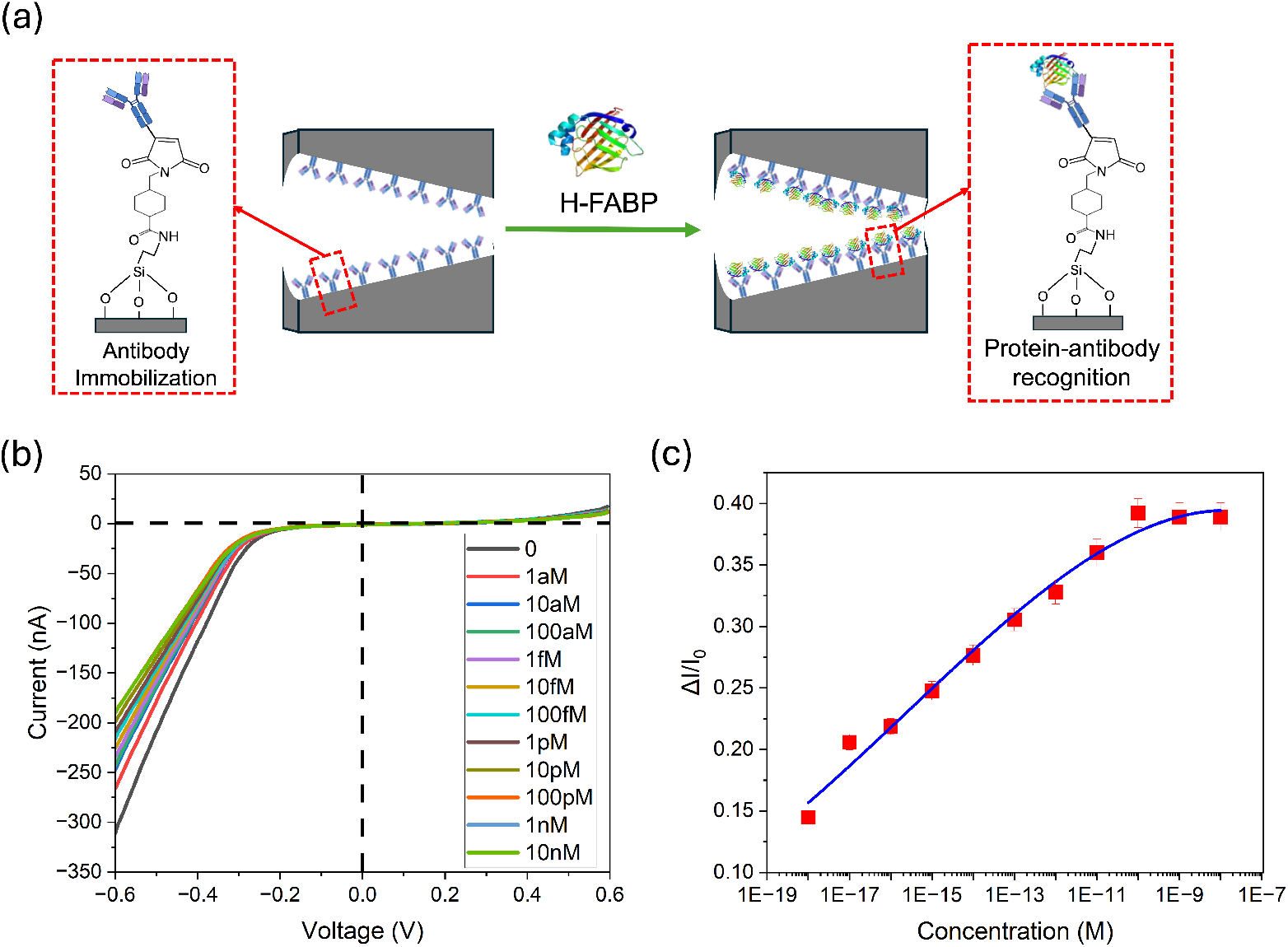
Detection of H-FABP using antibody-functionalized conical SiO_2_ nanopores. (a) Schematic illustration of specific recognition of H-FABP at the antibody immobilized nanopore surface. (b) Representative current-voltage (I-V) characteristics recorded after exposure to increasing concentrations of H-FABP, showing a concentration-dependent modulation of the ionic current. (c) Normalized current change (ΔI/I_0_) as a function of H-FABP concentration, demonstrating a clear binding-dependent response over a wide dynamic range (the solid line represents a fit to a binding isotherm).

**Figure 3:**
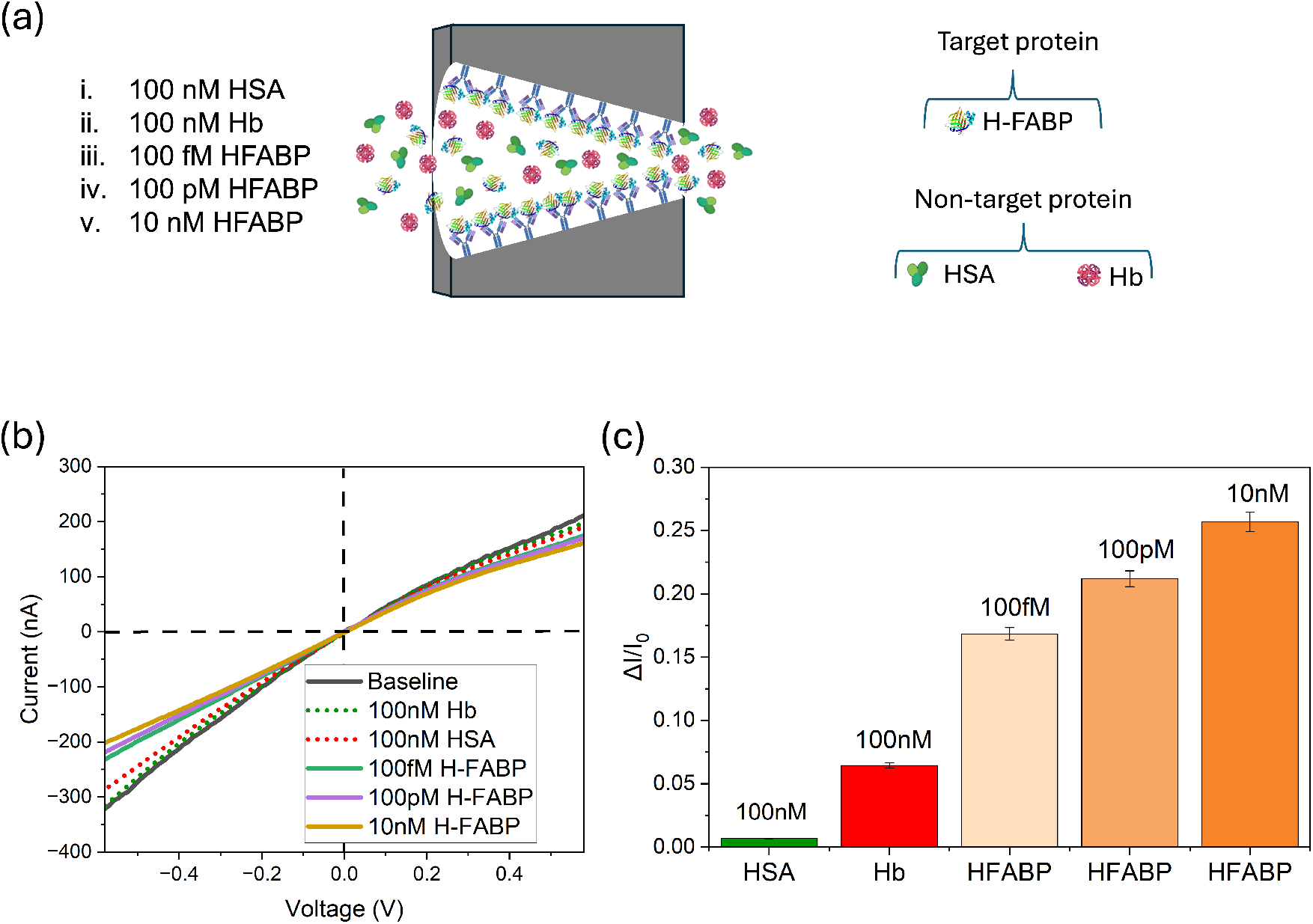
Selectivity of the antibody-functionalized conical SiO_2_ nanopore toward H-FABP. (a) Schematic illustration of the selectivity experiment using non-target proteins (HSA and Hb) and the target protein H-FABP at different concentrations. (b) Current-voltage (I-V) curves of H-FABP antibody modified conical nanopore in 10 mM NaCl-PBS (pH7.2) solution with 100 nM HSA, 100 nM Hb, and increasing concentrations of H-FABP, showing only small current modulation for non-target proteins and a concentration-dependent response for H-FABP. (c) Corresponding normalized current changes (ΔI/I_0_), highlighting the high specificity of the biosensor toward H-FABP over non-specific protein interactions.

**Figure 4:**
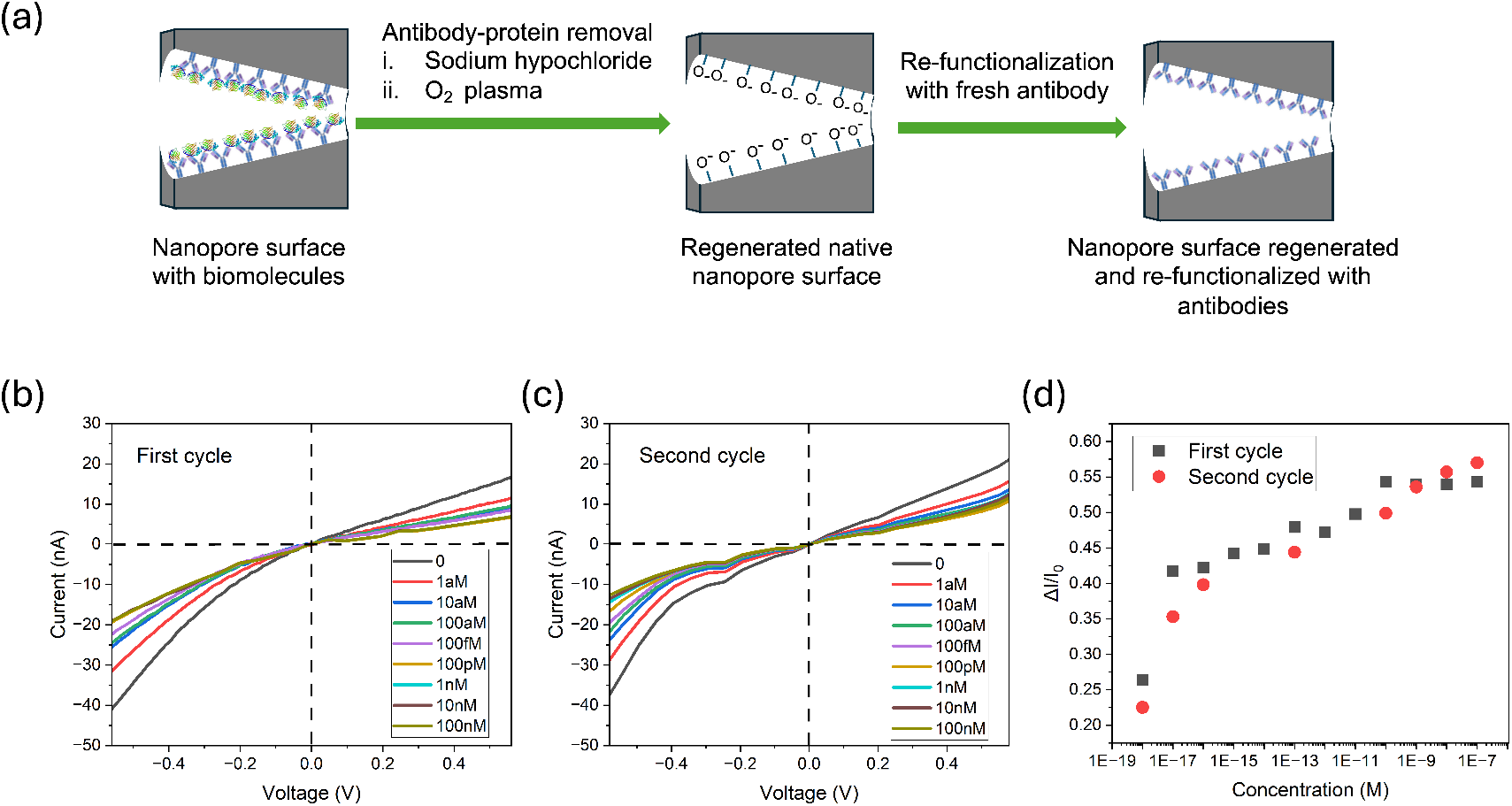
Regeneration and reusability of the antibody-functionalized conical SiO_2_ nanopore sensor. (a) Schematic illustration of the regeneration cycle, in which bound biomolecules are removed to restore the native nanopore surface charge, followed by re-functionalization with new antibodies, enabling reuse of the nanopore membrane. (b) Current-voltage (I-V) curves recorded during the first sensing cycle after exposure to increasing concentrations of the target protein. (c) Current-voltage (I-V) curves obtained during the second sensing cycle after regeneration, demonstrating reproducible rectification behavior and signal response. (d) Normalized current change (ΔI/I_0_) as a function of analyte concentration for the first and second cycles, confirming reproducible sensor performance and effective regeneration without loss of sensitivity.

Figure 1b shows a schematic of the SiO_2_ nanopore surface functionalization process. The native nanopore carries a negatively charged silanol group at physiological pH (pH 7.2), which generates a characteristic rectifying current-voltage (I-V) curve. Exposing the nanopore to the APTES vapor introduces amine functionalities on the SiO_2_ surface that carry a positive charge at the neutral pH range. The subsequent reaction with a sulfo-SMCC crosslinker solution generates a maleimide-acctivated surface, enabling covalent immobilization of the H-FABP antibody onto the pore walls. These sequential modifications lead to distinct changes in the ionic current response, allowing each functionalization step to be directly monitored through its characteristic shift in the I–V curve and the corresponding ion current rectification (ICR), shown in Figure 1c and d, respectively. The magnitude of ionic current rectification (ICR) can be quantitatively determined from the logarithmic ratio of the current measured at a given voltage polarity to the current measured under an equal voltage magnitude but opposite polarity.^23,24^ Functionalization of the nanopore surface with aminosilane alters the rectification ratio from (-0.13 ± 0.1) to (+0.98 ± 0.03), and then to (-1.26 ± 0.04) after modification with the antibody. These distinct and reproducible changes in the rectification ratio confirm the successful assembly of the respective chemical and biomolecular layer on the nanopore surface.

After preparing the antibody-modified nanopore membrane, we evaluated its sensing performance for H-FABP by measuring the current–voltage response of the system before and after introducing various concentrations of the protein (H-FABP) in 10 mM NaCl–PBS buffer at pH 7.2. The immobilized H-FABP antibodies on the pore walls selectively bind H-FABP protein molecules in the solution with high affinity. As a result, the antibody–protein interaction (Figure 2a) partially screened the negative surface charge of the nanopore walls, leading to a reduction in the ion current rectification ratio. Figure 2b shows the I-V curves for the H-FABP antibody-modified nanopore membrane after exposure to increasing concentrations of H-FABP protein. Over a broad concentration range between 1aM and 10nM, the ionic current measured at -0.6 V dropped considerably, whereas the current at +0.6 V remained largely unchanged. The absolute value of current change ratio, ΔI/I_0_ at –0.6 V is shown in Figure 2c. This value serves as a quantitative measure of how the ionic current varies with increasing protein concentrations. The ratio increases linearly for low protein concentrations over 8 orders of magnitude in a range between 1 aM and 100 pM. Beyond this range, the ratio plateaued and remained relatively unchanged as the concentration increased up to 10 nM. This behavior is likely due to saturation of H-FABP binding on the nanopore surface at higher concentrations. These results demonstrate ultra-high sensitivity of our biosensing platform to the target analyte, achieving a detection limit (LOD) as low as 0.4 aM. Details of the LOD calculation are provided in the Supporting Information. This detection limit for H-FABP is significantly lower than those reported for existing analytical methods, which typically achieve limits of detection in the pM range, with the lowest reported value being approximately 56 pM using a capacitive immunosensor.^12,18–20^

As an effective biosensing platform, the sensor must exhibit high selectivity toward its target analyte. Figure 3a presents a schematic illustration of the selectivity test performed using both target proteins (H-FABP) and non-target proteins human serum albumin (HSA) and hemoglobin (Hb) passing through the nanopore. For this test, we first recorded the baseline I–V curve of the antibody-modified nanopore in 10 mM NaCl–PBS buffer. Subsequently, 100 nM HSA and 100 nM (Hb) solutions were introduced, followed by increasing concentrations of H-FABP, the lowest 6 orders of magnitude lower than those of the non binding proteins. Figure 3b shows the corresponding I–V plots. Both 100 nM HSA and 100 nM Hb produced negligible changes in the I–V curves, indicating a lack of binding to surface-bound antibodies as well as non-specific binding to the nanopore surface. In contrast, only 100 fM H-FABP generated a clear and measurable decrease in the ionic current, which further decreased with the introduction of 100 pM and 10 nM H-FABP, highlighting the fact that the distinct signal is produced only by the target protein. We also determined the ratio between the change in ionic current at –0.6 V from the I-V plots and summarized the results in Figure 3c. It is apparent that the translocation of HSA and Hb through the functionalized nanopore only resulted in small changes in the ionic current, while the introduction of 100 fM, 100 pM, and 10 nM H-FABP produced significant changes. These results are consistent with the specific binding of the target protein to the immobilized antibodies and demonstrate excellent bioselectivity *>*6 orders of magnitude, clearly distinguishing the target protein from other non-specific proteins.

Regeneration experiments were performed to evaluate the stability and reusability of our nanopore membranes and the reproducibility of the sensing performance, which are essential characteristics for practical biosensing applications. After target detection, antibody-functionalized nanopores were regenerated using sodium hypochlorite treatment followed by O_2_ plasma cleaning for 5 minutes (gas flow rate of 10 sccm), and subsequently refunctionalized with fresh antibodies. Figure 4a shows the schematic illustration of regeneration and reuse. Figure 4b shows the current-voltage plots of the first sensing cycle. One can see a clear concentration dependence of the ionic current with increasing H-FABP concentration, similar to that presented in Fig. 2b. After regeneration and re-functionalization, the nanopore demonstrated a comparable response in the second cycle (shown in Figure 4c), with similar shifts in the current voltage curves observed across the same concentration range. This suggests successful re-functionalization of the nanopores with the H-FABP antibody to a similar level. In addition, the corresponding relative current changes ΔI/I_0_ at -0.6 V, calculated from the Figures 4b and 4c are shown in Figure 4d. In both cases, ΔI/I_0_ increased steadily with increasing H-FABP concentration at a very similar rate. These experiments confirm that nanopore membranes are reusable, and regeneration leads to a comparable sensing performance.

## Conclusion

In this work, we have presented an ultrasensitive, highly selective, and reusable platform for biomolecular sensing using conical SiO_2_ nanopores. The selectivity of *>*10^6^ is determined by the specific recognition between the target proteins and the antibodies covalently immobilized on the surface of the nanopore. The selective binding of the proteins induces changes in the nanopore surface charge that lead to an ultra-low detection limit in the aM range for the given membrane and pore system. The sensing performance was demonstrated for the detection of H-FABP, which is an important biomarker of cardiac injury and is also emerging as a promising clinically relevant biomarker in neurodegenerative disorders.^11,17^ The limit of detection of our nanopore-based biosensor of approximately 0.4 aM for H-FABP, is several orders of magnitude lower than that of existing sensing techniques.^12,18–20^ The biomolecular immobilization strategy demonstrated in this experiment is broadly applicable and can be extended to other biomarkers, e.g., we also demonstrated sensing of bovine serum albumin (BSA) with similar sensitivity and selectivity (see supplementary information). The measurement process only requires several minutes and yields quick results important for assessment of cardiac diseases. Although clinical cutoff concentrations for H-FABP in serum are in the range of 2–6 ng/mL (approximately 130–400 pM), the ability to sense H-FABP at ultralow concentrations is particularly important for neurodegenerative biomarker applications, where protein levels in blood or saliva can be substantially lower, in particular during the early stages of disease. In addition, such extreme sensitivity enables extensive sample dilution (*>*10^5^-fold), effectively suppressing matrix effects and minimizing pore clogging that commonly limit nanopore analysis of undiluted physiological fluids.^5^ Overall, this work highlights the potential of conical nanopores in SiO_2_ as a robust biosensing platform for early-stage biomarker detection and lays the foundation for extending this approach to other clinically relevant targets.

## Experimental

The nanopores were fabricated in free standing SiO_2_ windows with a thickness of (930 ± 5) nm and dimension of 50 *µ*m × 50 *µ*m, supported by 5.6 mm × 5.6 mm silicon frames. The windows were prepared using standard MEMS techniques that included RCA cleaning, photolithographic patterning, backside SiO_2_ removal by reactive ion etching, and anisotropic silicon wet etching. ^25^ Nanopores were fabricated using the ion-track etching technique, which involves high energy ion irradiation followed by chemical etching. A detailed description of these processes are given in our previous work.^8,9^ The etching time required to achieve the desired nanopore tip diameter was determined using our previously developed etching model,^26^ resulting in an optimized etching time of 21 minutes. The total number of nanopores formed in each membrane was estimated from SEM image analysis. In these experiments, three independently fabricated membranes were used containing approximately 25, 23 and 43 nanopores, respectively. The tip diameter of the conical nanopores was determined using conductometric measurements carried out in a custom-built setup consisting of two half-compartments. Both compartments were filled with a 10 mM NaCl solution prepared in phosphate-buffered saline (PBS). Sodium chloride and BupH™ Phosphate Buffered Saline Packs were purchased from Thermo Fisher Scientific (Product No. 207790010 and 28372, respectively). Given that the isoelectric point of the SiO_2_ nanopore system is 4.5 ± 0.1,^9^ all measurements were performed at near neutral pH conditions (pH 7.2) to ensure a negatively charged pore surface.

The pH and bulk conductivity of the electrolyte solutions were measured using a Thermo Fisher Scientific Orion Star^*T M*^ A215 tabletop multiparameter meter. For conductometric measurement, two Ag/AgCl electrodes were placed in the electrolyte-filled compartments and connected to a source meter (Keithley 2450). An applied voltage, *V* across the membrane generated an ionic current, *I*, which was recorded simultaneously. The conductance deduced from the current-voltage plot is used to determine the nanopore tip radius.^8,9^ Since the I-V characteristics are non-linear at 10 mM NaCl, the conductance was extracted from the linear region of the I–V plot between -0.1 V and +0.1 V.

The antibody immobilization process involves four main steps. First, the nanopore membranes were washed with deionized water and then air-dried. The membranes were treated with oxygen plasma for 5 minutes to introduce negative hydroxyl groups on the nanopores surface using a Tergeo plasma cleaner obtained from Pie Scientific (RF power of 25 W and gas flow rate of 10 sccm) and subsequently baked at 110^*°*^C for 20 minutes. For vapor-phase silanization, 0.5 ml of 3-Aminopropyl triethoxysilane (APTES, Sigma-Aldrich, Cas No. 919-30-2) was placed in a 250 mL desiccator together with the membranes. The chamber was evacuated to reduce ambient moisture and silanization was carried out at an elevated temperature (typically 80 to 85 ^*°*^C) for approximately 2 h. The membranes were then rinsed with anhydrous ethanol and cured at 110^*°*^C for 20 minutes. A detailed description of these processes is given in the Supporting Information.

In the second step, the crosslinker solution was prepared by dissolving 2 mg of Sulfo-SMCC in 1 ml of coupling buffer (PBS-EDTA: 50 mM phosphate, 0.15 M NaCl, 10 mM EDTA, pH 7.2). This solution was immediately used to prevent hydrolysis. The silylated nanopore surface was then covered with the crosslinker solution to ensure a uniform coating. The nanopore membranes were incubated for 2 h at room temperature (∼25 ^*°*^C) to allow the reaction to proceed. Following incubation, the modified nanopore surface was thoroughly rinsed with the coupling buffer solution to remove any unbound reagent.

In the third step, the antibody was modified and activated with sulfhydryl groups to enable binding to the maleimide-activated amino-modified nanopore surface. First 25 *µ*l of antibody was dissolved in 475 *µ*l of pH-adjusted coupling buffer (pH8). Separately, 2 mg of Traut’s Reagent (2-iminothiolane·HCl) was dissolved in 1 ml of the same buffer. Traut’s reagent was purchased from Thermo Fisher Scientific (Product No. 26101). Immediately after preparation, 25*µ*l of the Traut’s Reagent solution was added to the antibody solution. The mixture was incubated for 45 minutes at room temperature to allow thiolation. After incubation, the modified antibody was purified to remove excess Traut’s Reagent using a pre-equilibrated desalting column (Thermo Fisher Scientific, D-Salt^*TM*^ Dextran Desalting Columns, Product No. 43230) with a coupling buffer at pH 7.2 and 500*µ*l fractions were collected. The fractions containing the antibody were identified by measuring the absorbance at 280 nm using the Thermo Fisher Scientific Varioskan LUX Multimode Microplate Reader, and those that showed the highest absorbance peak were pooled. The resulting antibody, which now contains sulfhydryl groups, was used immediately in the subsequent step (more details in the Supporting Information). The H-FABP antibody was purchased from Neo-Biotechnologies (Cat. No. 2170-MSM8-p1ABX), and the corresponding recombinant protein was purchased from Creative BioMart (Cat. No. 6927H).

In the last step, the maleimide-activated nanopore membranes were covered with sulfhydryl-modified antibody solution. Nanopore membranes were then incubated for 4 h at room temperature (∼25^*°*^C) to allow covalent attachment of the antibody. Once incubation was complete, the reaction solution containing any unbound antibody was removed. The nanopore membrane was then thoroughly rinsed with coupling buffer solution to ensure that only covalently bound antibody molecules remained. A detailed illustration of this process is available in the Supporting Information.

The current–voltage characteristics of the antibody functionalized SiO_2_ conical nanopores were measured in 10 mM NaCl–PBS buffer (pH 7.2). The ionic current was recorded as the applied voltage was swept from −0.6 V to +0.6 V at a rate of 0.1 V s^*−*1^. This voltage window was selected to minimize the risk of protein denaturation that can occur at higher applied potentials. The error bars shown in each figure correspond to the measurement uncertainty obtained from at least three independent measurements performed on the same nanopore. All experiments were conducted at room temperature. Protein solutions (H-FABP) of various concentrations were prepared in 10 mM NaCl–PBS (pH 7.2) and used for current-voltage measurements.

After sensing experiments, antibody-functionalized nanopores were dipped in 1M sodium hypochlorite solution for 2 h followed by rinsing in deionized water for 15 minutes. The nanopores were then treated with O_2_ plasma for 5 minutes to remove residual organic species and regenerate the surface. Subsequently, the nanopore memrbanes were re-functionalized with antibodies using the previously described functionalization process.

## Acknowledgement

N.A. gratefully acknowledges the Australian National University for providing a URS scholarship. Part of the research was undertaken at the SAXS/WAXS beamline at the Australian Synchrotron, part of ANSTO, and the authors thank the beamline scientists for their technical assistance. This work used the ACT node of the NCRIS-enabled Australian National Fabrication Facility (ANFF-ACT). Additionally, we acknowledge the GSI Helmholtz Centre for Heavy Ion Research (GSI Helmholtzzentrum für Schwerionenforschung) in Darmstadt for their assistance with the ion irradiation experiments at the UNILAC accelerator X0 beamline (FAIR Phase 0).

