## Supplementary Information for "Detection of attomolar concentration of heart-type fatty acid binding protein using ion current rectification sensing with conical SiO_2_ nanopores"

### Characterization of conical nanopores in $SiO_2$

The morphology and dimensions of the $SiO_2$ nanopore membranes were characterized using scanning electron microscopy (SEM). Figures S1a-c show the plan-view SEM images of the membranes used in this study. Pore density and the total number of nanopores were determined individually for each membrane by direct counting from the plan-view SEM images. The pore density was estimated to be approximately 1 $\times 10^6$ pores cm$^{-2}$, corresponding to $\sim$25 nanopores within a 50 $\times$ 50 $\mu$m$^2$ window for representative samples. The three membranes used in this work contained approximately 25, 23 and 43 nanopores, respectively.

Higher-magnification plan-view images (Figure S1d) were used to determine the pore base diameter. Due to local charging at the pore edges, visible as a bright ring around each opening, the pore boundary was defined at the outer edge of the charged region. Using this approach, the average pore base diameter was determined to be (380 $\pm$ 5) nm. Cross-sectional SEM imaging (Figure S1e) confirmed that the pores possess a conical geometry that extends through the thickness of the membrane.

### Vapor-phase aminosilane functionalization

Vapor-phase silanization was carried out in a 250 mL sealed desiccator. Prior to silanization, the $SiO_2$ membrane was cleaned by $O_2$ plasma treatment for 5 min to remove organic contaminants and generate a hydroxylated surface. The membrane was then baked at 120 °C for 20 min to dehydrate the surface and promote the subsequent aminosilane attachment.

After dehydration, the membrane was exposed to APTES vapor. For this step, 0.5 mL of 3-aminopropyltriethoxysilane (APTES, Product No. 80370) was placed in a 40 mm glass Petri dish inside the desiccator. The cleaned and dehydrated membrane was positioned above the silane source, as shown in Figure S2. The chamber was evacuated to remove residual moisture and placed on a hot plate maintained at 80-85 °C. Vapor-phase silanization was allowed to proceed for 2 h.

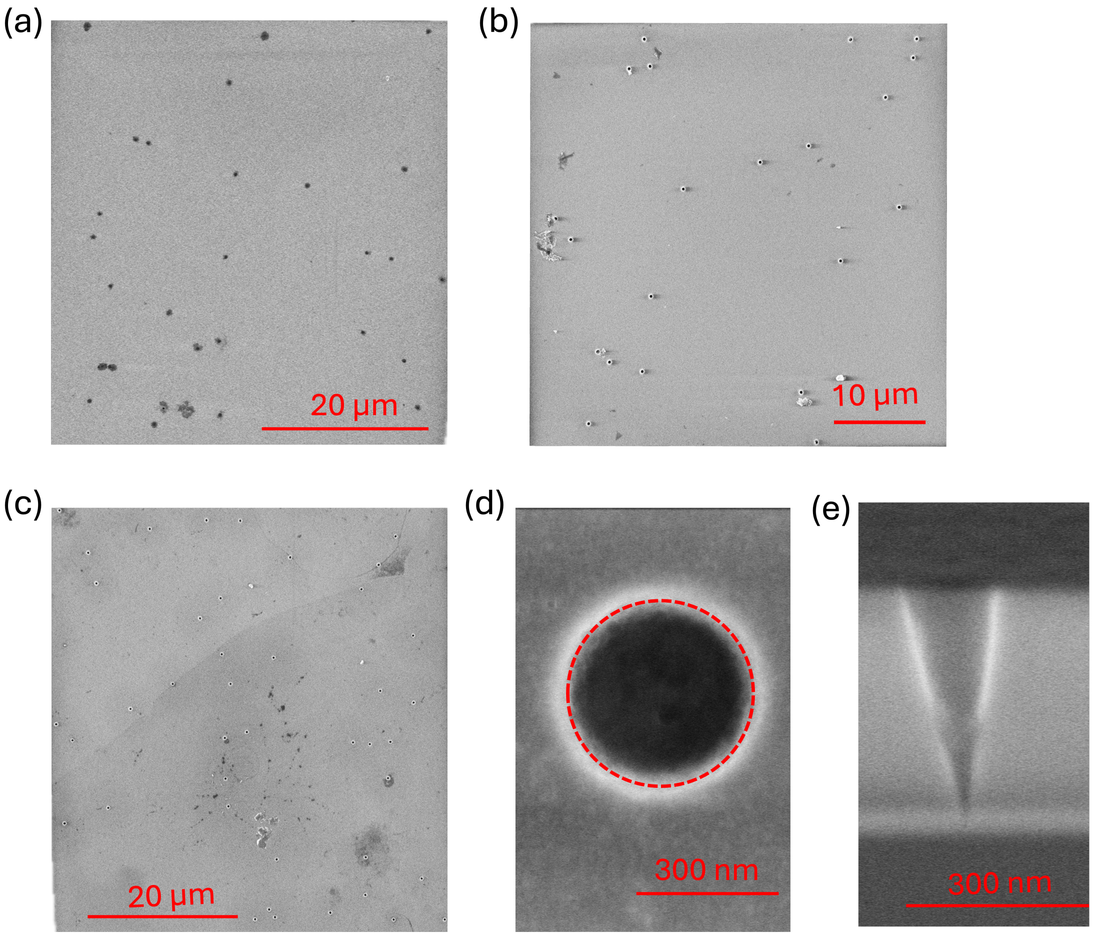


Figure S1: Morphological characterization of conical $SiO_2$ nanopore membranes. (a–c) Plan-view SEM images of the experimental membranes fabricated under identical irradiation and etching conditions, showing approximately 25, 23, and 43 nanopores within the active window for the respective membranes. (d) High-magnification plan-view image of a pore used to determine the pore base diameter. (e) Cross-sectional SEM image of nanopore confirming the conical pore geometry.

Following silanization, the membrane was removed from the chamber and rinsed with anhydrous ethanol to remove the unreacted silane species. The membrane was then baked at 110 °C for an additional 20 min to promote the formation of siloxane bonds and remove weakly bound aminosilane groups. Successful silanization was verified by changes in the current–voltage characteristics of the nanopores.

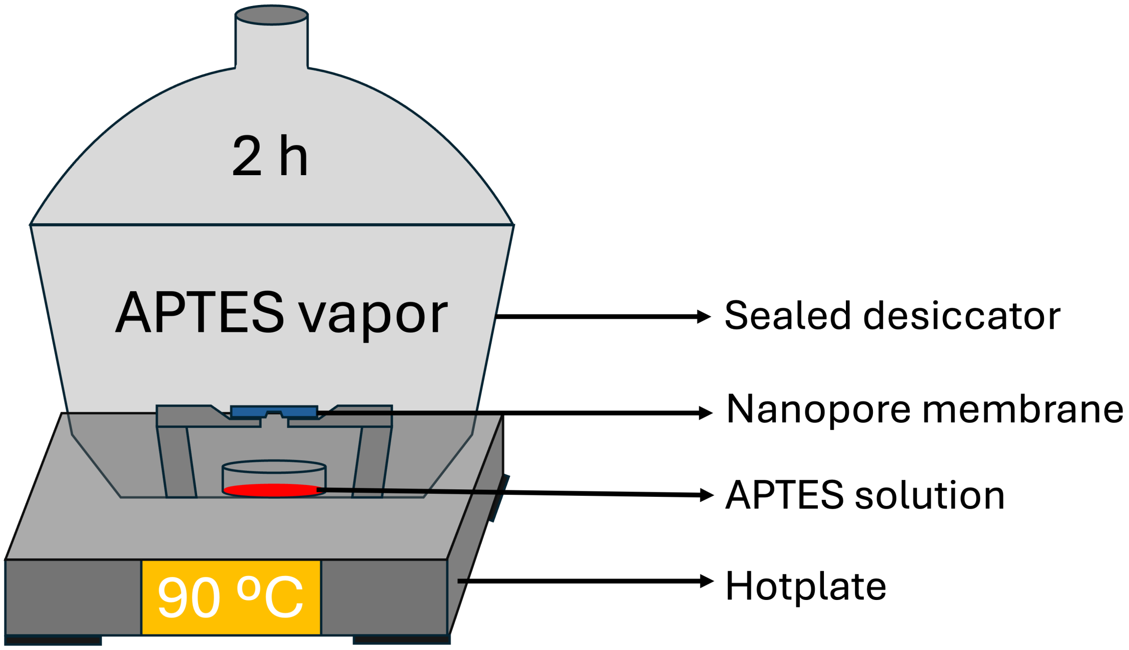


Figure S2: Surface silanization process of nanopores $SiO_2$ membranes.

#### Functionalisation of the nanopore membrane

Figure S3 schematically illustrates the antibody activation procedure and the subsequent fabrication of the nanopore biosensor. Antibodies were first activated using Traut's reagent to introduce reactive thiol groups. The reaction mixture was vortexed and incubated at 26 °C for 45 min, followed by purification using a desalting column to remove excess reagent. The activated antibodies were then quantified and isolated using a microplate reader prior to immobilization. The whole procedure is explained in the Figure S3a.

Bio-functionalization of the nanopore membrane is shown in Figure S3b. In this process,

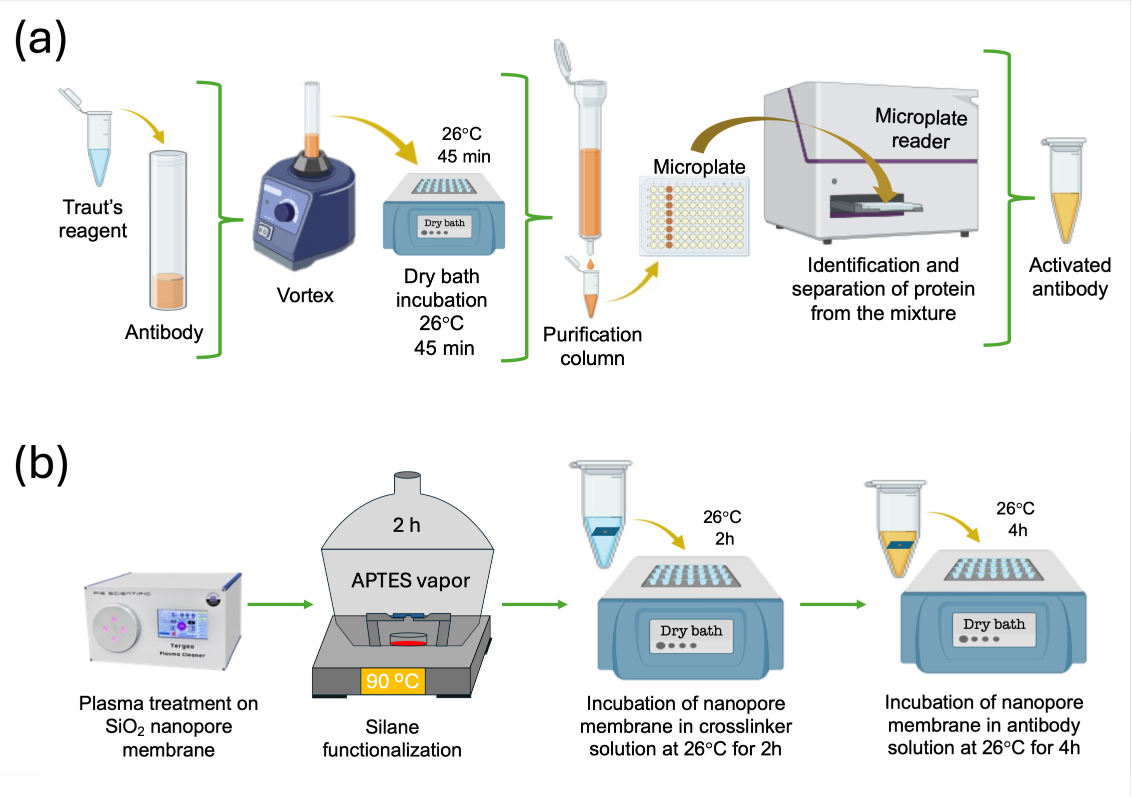


Figure S3: Schematic illustration of the antibody activation and bio-functionalization of the nanopore membrane. Antibody activation involving reagent mixing, incubation, and purification steps prior to immobilization, with activation efficiency assessed using a microplate reader (top). Functionalization of the nanopore membrane, including $O_2$ plasma cleaning, vapor-phase APTES silanization at 90 °C, and subsequent thermal curing steps to prepare the nanopore surface for antibody immobilization (bottom).

the $SiO_2$ nanopore membranes were initially cleaned and activated by oxygen plasma treatment to generate surface hydroxyl groups. The membranes were then functionalised with APTES via vapor-phase silanisation to introduce amine groups on the nanopore surface. After silanisation, the membranes were incubated in a crosslinker solution at 26 °C for 2 h to generate reactive surface groups for antibody attachment. Finally, the activated antibodies were immobilised by incubating the nanopore membranes in the antibody solution at 26 °C for 4 h, resulting in a stable antibody-functionalised nanopore biosensor.

#### Validation of the sensing performance of the nanopore-based biosensor

To validate the nanopore sensing platform, initial experiments were carried out using a common protein-antibody system based on bovine serum albumin (BSA). The nanopores were functionalised with BSA antibodies following the same surface-modification protocol described in the main text, and the sensing response toward BSA was evaluated over a wide concentration range. Figure S4a shows the current–voltage (I–V) plots of the BSA antibody-functionalised conical $SiO_2$ nanopores after exposure to increasing concentrations of BSA, ranging from 1 aM to 10 nM. The baseline response was measured with 10mM Nal-PBS buffer solution before the protein introduction. Concentration-dependent changes in the I–V curves are observed relative to the baseline, indicating successful binding of BSA within the nanopore and corresponding modulation of the ionic current. Figure S4b presents the normalized current change ($\Delta I/I_0$) extracted from the I-V data at -0.6 V as a function of BSA concentration. The signal increases monotonically with concentration and approaches saturation at higher concentrations.

Our current change data are well fit by a sigmoidal concentration–response curve exhibiting clear saturation at higher analyte concentrations. This behavior is characteristic of surface-binding-limited sensing and is well described by a Langmuir-type adsorption model, as commonly reported for nanopore and surface-immobilized biosensors.[1–3] The observed response is consistent with a binding-based sensing mechanism governed by antibody–protein interactions at the nanopore surface. Based on this validated sensing strategy, the platform was subsequently extended to the detection of H-FABP. Together, these results demonstrate that the sensor can be readily adapted to other protein targets through appropriate antibody functionalization.

Figure S4(c-d) evaluates the specificity and selectivity of the BSA antibody-functionalized nanopore membrane by comparing its response to BSA protein and a non-target protein, hemoglobin. Figure S4c shows the current–voltage (I–V) characteristics measured without protein (baseline), after exposure to the non-target protein (100 nM hemoglobin), and subsequently after introducing the target protein (BSA) at concentrations ranging from 100 fM to 10 nM. While BSA induces pronounced, concentration-dependent changes in the I–V response, hemoglobin results in only a minor deviation from the baseline, indicating weak nonspecific interaction with the functionalized nanopore surface. The corresponding normalized current change ($\Delta I/I_0$) extracted from the I–V data is shown in Figure S4d. At concentrations six orders of magnitude higher than the minimum concentration of BSA measured, hemoglobin yields a significantly smaller $\Delta I/I_0$ compared to BSA, while increasing BSA concentration results in a systematic increase in signal magnitude. These results confirm that the observed electrical response is predominantly from specific antibody–protein binding and demonstrate the selectivity of the nanopore sensing platform.

### Determination of limit of detection (LOD)

#### Limit of detection (LOD)

The limit of detection (LOD) was determined using the $3\sigma$ criterion in accordance with IUPAC recommendations.[4,5] The LOD in the signal domain was defined as the mean blank response plus three times the standard deviation of the blank signal. The concentration

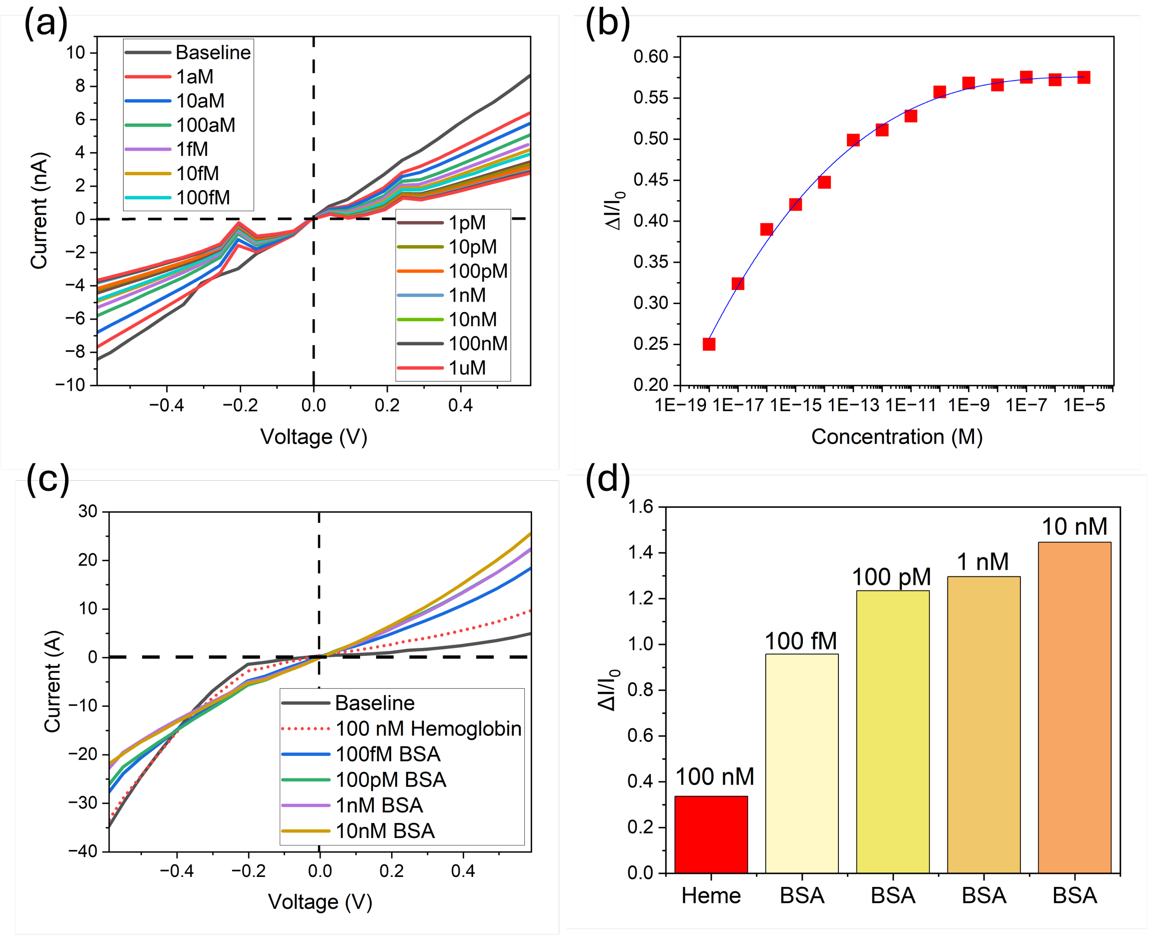


Figure S4: **Sensitivity and selectivity of the BSA antibody-functionalized nanopore membrane.** (a) Representative current-voltage (I-V) characteristics recorded after exposure to increasing concentrations of BSA, showing a concentration-dependent modulation of the ionic current. (b) Normalized current change ($\Delta I/I_0$) as a function of BSA concentration, demonstrating a clear binding-dependent response over a wide dynamic range (the solid line represents a fit to a binding isotherm). (c) Current-voltage (I-V) curves of BSA antibody modified conical nanopore in 10 mM NaCl-PBS (pH7.2) solution with 100 nM hemoglobin, and increasing concentrations of BSA, showing only small current modulation for non-target proteins and a concentration-dependent response for BSA. (d) Corresponding normalized current changes ($\Delta I/I_0$), highlighting the high specificity of the biosensor toward BSA over non-specific protein interactions.

LOD was obtained by converting the signal LOD using the calibration curve slope in the low-concentration regime. The sensor response was defined as

$$y = \frac{\Delta I}{I_0} = \frac{I_0 - I}{I_0}, \tag{1}$$

where $I_0$ is the baseline ionic current (blank) and $I$ is the current after exposure to analyte.

The LOD in the response domain was calculated as

$$y_{\mathrm{LOD}} = \bar{y}_{\mathrm{blank}} + 3\sigma_{\mathrm{blank}}, \tag{2}$$

where $\bar{y}_{\mathrm{blank}}$ and $\sigma_{\mathrm{blank}}$ are the mean and standard deviation of the blank response, respectively. Because the response was baseline-corrected, $\bar{y}_{\mathrm{blank}} \approx 0$, giving $y_{\mathrm{LOD}} = 3\sigma_{\mathrm{blank}}$.

The blank noise in $y$ was obtained by propagating the current noise:

$$\sigma_{\mathrm{blank}} = \sigma_y \approx \frac{\sigma_I}{|I_0|}, \tag{3}$$

where $\sigma_I$ is the standard deviation of the baseline current. Using $|I_0| = 3.10 \times 10^{-7}$ A and $\sigma_I = 6.6 \times 10^{-9}$ A, we obtained $\sigma_y = 2.13 \times 10^{-2}$ and thus $y_{\mathrm{LOD}} = 6.39 \times 10^{-2} \approx 0.063$.

The concentration LOD was extracted from the calibration curve $y = f(C)$ by inversion:

$$C_{\mathrm{LOD}} = f^{-1}(y_{\mathrm{LOD}}). \tag{4}$$

For a low-concentration extrapolation using the lowest calibration point ($y = 0.1447$ at $C = 1.0 \times 10^{-18}$ M), this yields

$$C_{\mathrm{LOD}} \approx 1.0 \times 10^{-18} \times \frac{0.063}{0.1447} = 4.35 \times 10^{-19} \text{ M } (\approx 0.43 \text{ aM}), \tag{5}$$

noting that this value is extrapolated below the lowest measured concentration.

### Regenaration and reproducibility of the nanopore-based biosensor

The reproducibility of the nanopore sensing platform was evaluated using the BSA antibody–protein system as an independent validation experiment. Two consecutive sensing cycles were performed with the conical $SiO_2$ nanopore functionalized with BSA antibody under identical experimental conditions. Figures S5a and S5b show the current–voltage characteristics recorded during the first and second sensing cycles, respectively, after exposure to increasing concentrations of BSA together with the corresponding baseline measurements. In both cycles, systematic and concentration-dependent modulation of the I-V response is observed.

Figure S5c compares the normalized current change ($\Delta I/I_O$) as a function of BSA concentration extracted from the first and second sensing cycles. The close agreement between the two datasets across the entire concentration range demonstrates good reproducibility of the sensing response and indicates that the antibody-functionalized nanopore remains stable during repeated measurements. These results confirm the robustness of the sensing strategy and support the applicability of our memrbrane system for reliable protein detection.

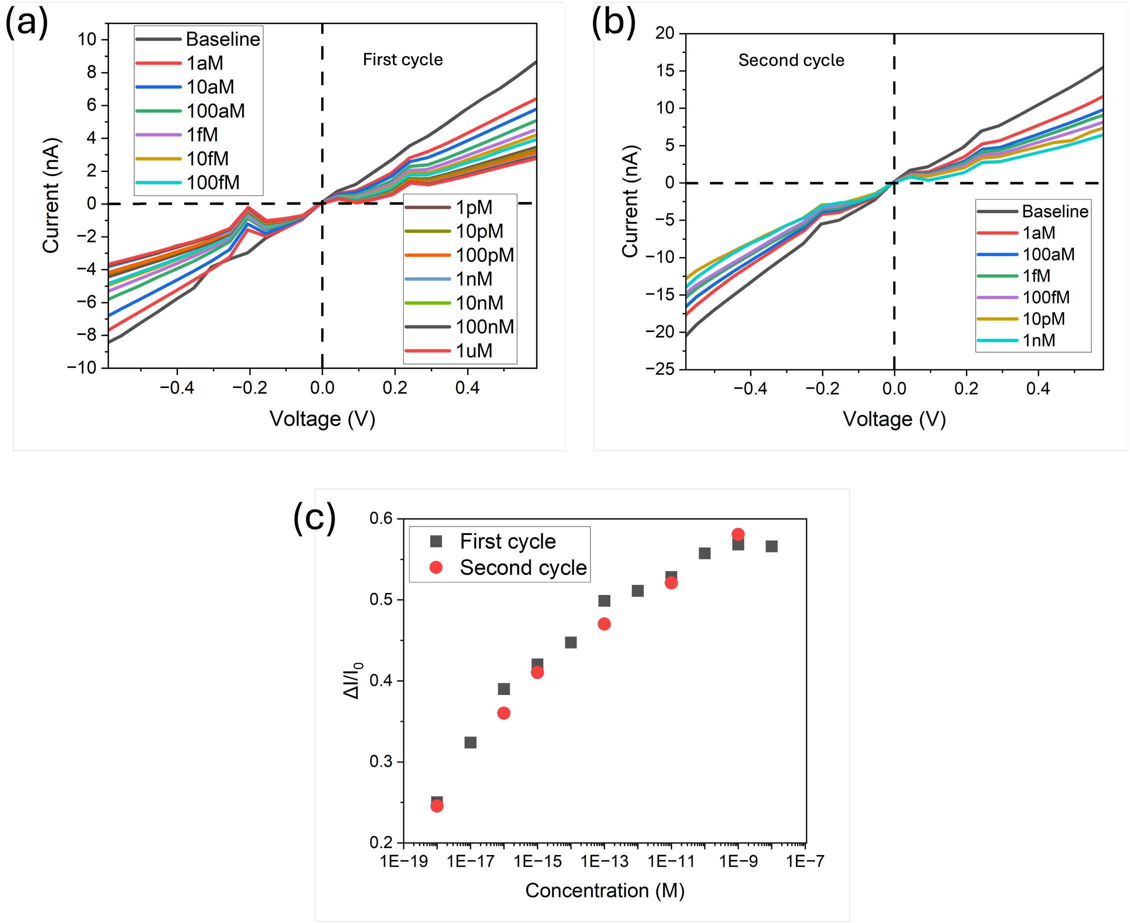


Figure S5: **Regeneration and reusability of the BSA antibody-functionalized conical $SiO_2$ nanopore sensor.** (a) Current-voltage (I-V) curves recorded during the first sensing cycle after exposure to increasing concentrations of the target protein. (b) Current-voltage (I-V) curves obtained during the second sensing cycle after regeneration, demonstrating reproducible rectification behavior and signal response. (c) Normalized current change ($\Delta I/I_0$) as a function of analyte concentration for the first and second cycles, confirming reproducible sensor performance and effective regeneration without loss of sensitivity.
